# Functional Divergence of Axon-Carrying Dendrite (AcD) and NonAcD Cells in Learning and Stability

**DOI:** 10.1101/2025.09.15.676360

**Authors:** Sima Hashemi, Shirin Shafiee, Christian Tetzlaff

**Affiliations:** III. Institute of Physics – Biophysics, Faculty of Physics, University of Göttingen, Göttingen, Germany; Group of Computational Synaptic Physiology, Department of Neuro- and Sensory Physiology, University Medical Center Göttingen, Göttingen, Germany

## Abstract

Axon-carrying dendrite (AcD) cells are a specialized class of hippocampal neurons where the axon initial segment originates from a basal dendrite rather than the soma, creating a privileged pathway for excitatory inputs on AcD branches to bypass perisomatic inhibition. However, their functional role in learning and synaptic stability remains unclear. To address this question, we modeled a CA3-CA1 network to compare the learning dynamics and synaptic stability of AcD and nonAcD cells. The results revealed that, during learning, these cell types employ distinct mechanisms. AcD cells primarily adopt a single-modal strategy, with all dendritic branches converging to encode inputs from a single assembly, whereas nonAcD cells follow a multi-modal approach, with individual branches encoding inputs from distinct assemblies. Additionally, consistent with experimental findings, our results suggest that during periods of high inhibition (such as ripples), AcD cells maintain stable synaptic weights, unlike the synaptic decay observed in nonAcD cells. These results, in line with experimental evidence, suggest that although the morphological distinction between AcD and nonAcD cells was long overlooked, it proves to be important, as it results in functional differences in learning mechanisms and in the capacity for stable information storage, highlighting their key role in learning and memory consolidation.

## Introduction

The canonical model of neurons involves the reception of inputs through dendrites, signal integration in the soma, and output transmission via the axon. However, in 2014, Thome et al. introduced a novel class of neurons in which the axon initial segment (AIS) originates from a basal dendrite, rather than the soma [31]. Figure 1 presents a schematic of both cell types. These axon-carrying dendrite (AcD) cells have been identified across multiple species and in several brain regions [30, 31, 35]. In non-primate mammals, these cells constitute approximately 10% to 21% of pyramidal neurons in cortical layers II–VI [35]. In mice, AcD cells are significantly more abundant in the CA1 region, comprising approximately 52.2% ± 2.2% of neurons, compared to the CA3 region at 28.3% ± 7.1% and the subiculum at 21.3% ± 3.7% [31]. Moreover, in CA1 region AcD cells predominantly occupy superficial positions [31].

**Figure 1:**
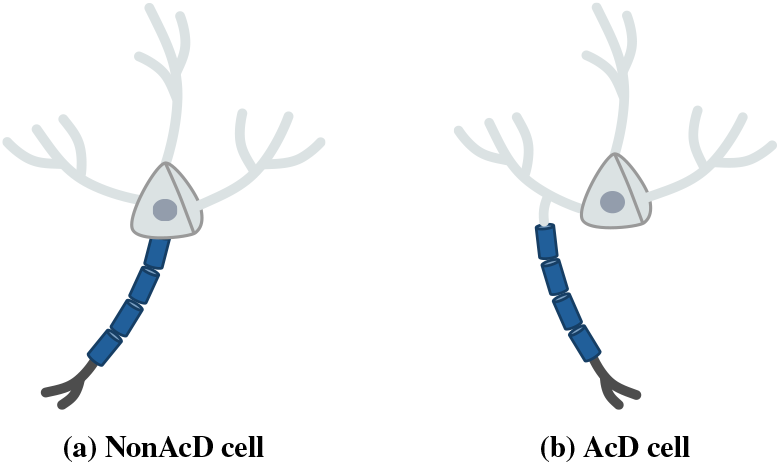
Schematic of AcD and nonAcD neurons. For both cells, the dendritic tree, soma, and axon are illustrated. In the AcD cell, the axon emerges from a basal dendrite rather than from the soma.

Morphological features are generally similar between AcD and nonAcD cells [17, 30]. Both pyramidal neuron types in the CA1 region typically possess three primary basal dendrites [17, 30]. Dendritic spine densities are also comparable between AcD and nonAcD neurons and remain consistent across AcD and nonAcD branches within individual AcD cells [30]. Although the overall basal dendritic trees of both cell types are similar in size and complexity, individual AcD branches are generally longer than nonAcD branches [30]. This greater branch length, combined with comparable spine density, results in a higher total number of spines on AcD branches. In line with this, it has been proposed that the greater number of spine–axon proximities between AcD branches and the contralateral CA3 region is the primary reason why AcD cells receive stronger excitatory input from that region compared to nonAcD cells [30]. Consequently, neurons with AcD morphology may be pivotal in facilitating interhemispheric interactions and serve as key hubs for both short-term and long-term changes in these connections [12, 26, 30].

Despite similarities in morphological features, AcD cells possess a privileged input pathway. The AcD branches exhibit increased intrinsic excitability, leading to a higher probability of dendritic spikes compared to nonAcD branches [31]. Furthermore, unlike typical neurons where dendritic inputs are subject to perisomatic inhibition before integration and output firing, AcD branches allow inputs to bypass this inhibition [17, 31]. As a result, these branches are potentially privileged pathways for directly coupling excitatory synaptic inputs to action potentials (APs) in many CA1 neurons [31]. This implies that during phases of strong inhibition, excitatory inputs are most efficient at the AcD branches [17, 31]. Correspondingly, experimental evidence demonstrates that during ripple oscillations in awake mice, AcD cells are far more likely to spike than nonAcD cells [17]. In summary, while AcD and nonAcD cells share similar dendritic structures, AcD cells function as privileged input sites, where excitatory inputs are more likely to generate dendritic spikes and efficiently trigger APs [17, 30, 31].

In this study, we aimed to investigate how the privileged input pathway affects learning and synaptic weight stability in AcD cells. To address this question, we simulated a simplified CA3-CA1 network, incorporating the experimentally identified differences between AcD and nonAcD cells. The model incorporates three key differences between AcD and nonAcD cells: 1) AcD cells have one AcD and two nonAcD dendritic branches, while nonAcD cells possess three nonAcD branches, consistent with experimental data on the count of basal dendritic branches [17, 30]; 2) the number of putative synapses (representing spines in neurons) on AcD branches is higher than that on nonAcD branches, in agreement with experimental evidence [30]; and 3) inputs to AcD branches bypass perisomatic inhibition before reaching the AIS compartment, unlike inputs to nonAcD branches. This aligns with experimental findings showing that stimulation of AcD branches generates action potentials with more hyperpolarized thresholds compared to nonAcD branches [31].

Using this model, we simulated both an Exploration and a Ripple phase to investigate learning dynamics and synaptic weight stability, respectively. Our results suggest that the privileged input pathway in AcD branches leads to distinct learning mechanisms in AcD and nonAcD cells, indicating that AcD and nonAcD cells may be specialized for different functions, with each playing a role in distinct aspects of information processing. Additionally, during periods of high inhibition, such as during ripple events, AcD cells exhibit greater stability in synaptic weights. In contrast, synaptic weights in nonAcD cells are less stable and more prone to forgetting. These findings are consistent with experimental data showing that CA1 pyramidal cells in the superficial layers—where AcD cells predominate—are more stable, whereas pyramidal cells in the deeper layers, which are primarily nonAcD cells, exhibit more hyperpolarized firing thresholds and are less stable during these events [14, 27, 31, 33]. This implies that AcD cells may be better suited for storing critical information than nonAcD cells, due to their higher stability.

## Materials and Methods

Here, we reconstructed a simplified CA3–CA1 network using a rewiring learning rule based on structural, functional, and backpropagating action potential plasticities. A schematic of this network is shown in Figure 2A.

**Figure 2:**
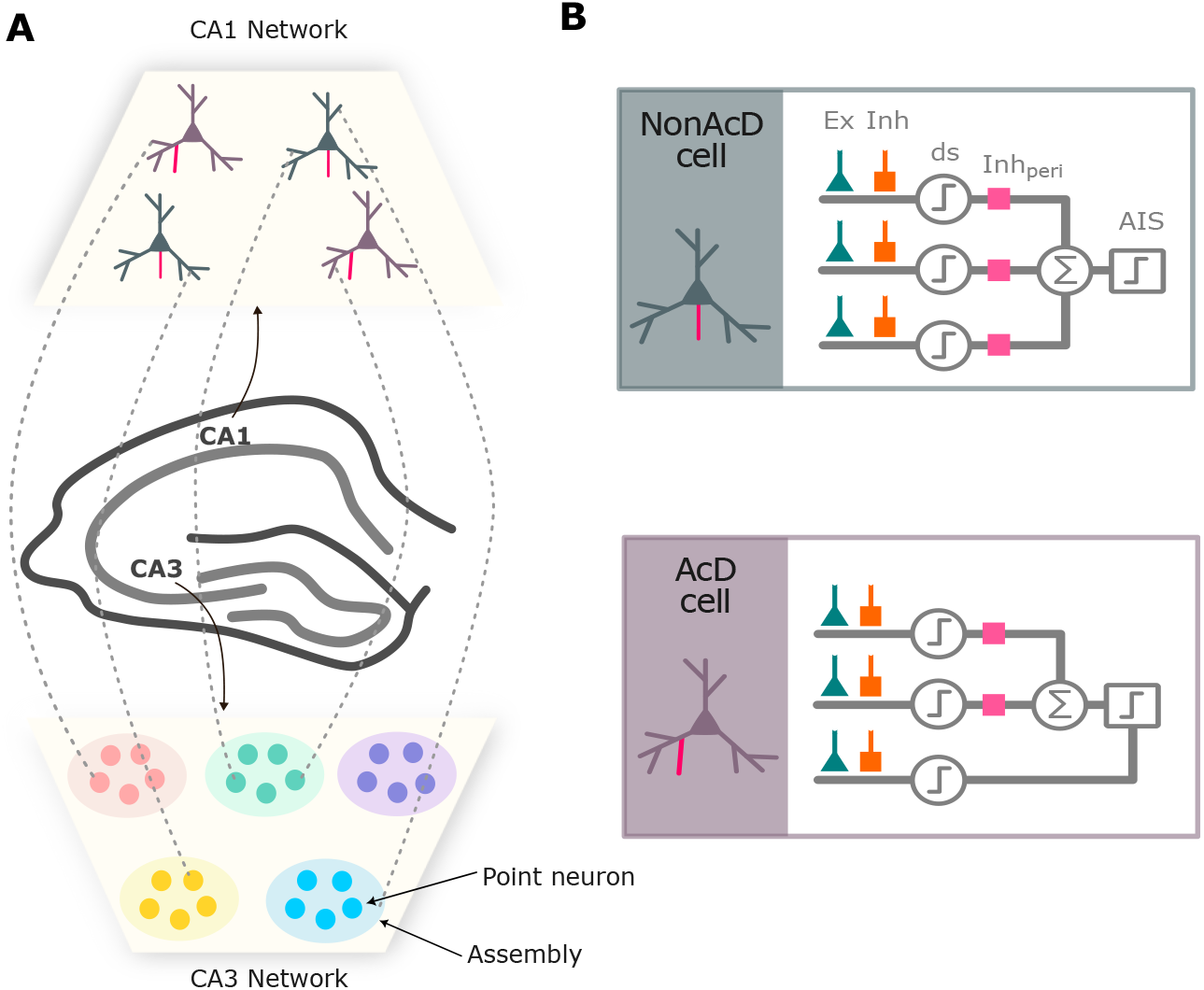
Schematic of the Reconstructed Model of the CA3-CA1 Network. **A)** *CA3-CA1 Network:* The CA3 network consists of 270 point neurons organized into 9 assemblies of 30 neurons each, while the CA1 network comprises 100 compartmental neurons, with 50% of their population designated as AcD cells, based on experimental data [31]. For simplicity, only the connections between CA3 and CA1 cells are assumed to be plastic, with recurrent connections within CA3 and CA1 networks omitted. **B)** *CA1 Compartmental Cell:* The schematic of AcD and nonAcD cells shows three basal dendrites, the soma, and the axon (pink). In our model, both cell types are represented by three dendritic compartments and a single AIS compartment. Each dendritic compartment receives excitatory input from CA3 neurons (green) and a direct inhibitory current (orange), representing a simplified abstraction of inhibitory signals from CA1 interneurons onto pyramidal cells. Integration of these inputs can generate a dendritic spike (d-spike) that propagates toward the AIS. In nonAcD branches, dendritic currents pass through perisomatic inhibition (pink box) before reaching the soma. The soma, depicted as a summation point, integrates these inputs and transmits positive currents to the AIS. By contrast, the privileged AcD branch bypasses perisomatic inhibition and connects directly to the AIS compartment.

### CA3–CA1 Network

The CA3 network was modeled with 270 point neurons organized into 9 distinct assemblies of 30 neurons each, while the CA1 network consisted of 100 compartmental neurons, 50% of which were designated as AcD cells, in accordance with experimental data [31]. For simplicity, we assumed that only the connections between CA3 and CA1 cells are plastic, whereas all recurrent connections within the CA1 and CA3 networks were neglected.

### CA1 Cells

In this model, we simplify the morphologies of AcD and nonAcD cells by representing each neuron as a compartmental model with three basal dendrites—each treated as a single compartment—and a separate AIS compartment, as illustrated in Figure 2B. Each dendritic compartment receives two types of input: excitatory input from CA3 neurons and an inhibitory direct current, representing a simplified abstraction of the connections between interneurons and pyramidal cells in CA1. A dendritic spike can be emerged on each branch when the values and timing of these two inputs satisfy voltage requirement for generation of dendritic spike. Subsequently, the generated dendritic spike propagates toward the AIS compartment. In nonAcD branches, dendritic spikes are subjected to perisomatic inhibition before they are integrated at the soma. In our model, soma is treated as an integrator unit of all incoming dendritic currents from nonAcD branches along with perisomatic inhibition, transmitting only the resultant net positive current to the AIS. However, the AcD branch connects directly to the AIS compartment, bypassing perisomatic inhibition entirely. This direct pathway emphasizes the unique role of the AcD branch in neuronal signaling.

### Dendritic Branches

The membrane potential of the dendritic branch *j* ∈ {1, 2, 3} at time *t* is represented by 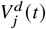. The dynamics of the membrane potential are defined as follows:

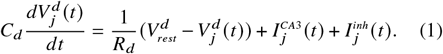

Here, *Cd, Rd*, and 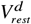 represent the dendritic membrane capacitance, resistance, and leak reversal potential, respectively. 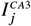 denotes the input current from the CA3 region to branch *j*. This input current is a weighted sum of all past spikes (represented by an alpha function) that occurred in each CA3 cell connected to this branch.

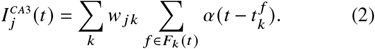

Here, *w*_*jk*_ represents the synaptic weight between dendritic branch *j* and CA3 input neuron *k*. The term *F*_*k*_ (*t*) denotes the set of all spikes of input neuron *k* that occurred before time *t*. The current from each connected input neuron is modeled as an alpha function with a synaptic time constant *τ*_*syn*_.

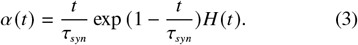

In addition to excitatory input from the CA3 region, CA1 cells receive inhibitory currents from interneurons. In general, two types of interneurons provide inhibitory signals to CA1 cells: parvalbumin-expressing interneurons (PVs) primarily target the perisomatic region, whereas somatostatin-expressing interneurons (SOMs) project to the dendritic tree [21, 34]. PVs target mainly the perisomatic region, while SOMs project to the dendritic tree [21, 34]. PV-mediated inhibition is fast and transient, whereas SOM-mediated inhibition is slowly recruited and persistent, increasing with high-frequency input [21, 34]. To simplify the model, instead of explicitly simulating inhibitory interneurons, we represent this inhibition as an external direct current added to each dendritic branch, denoted 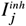, which could mimic the overall inhibitory influence of interneurons on the basal dendrites.

### Dendritic Spike

If the dendritic branch potential exceeds the spike threshold 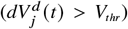, a dendritic spike is triggered, and the branch enters a refractory period of duration 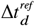. Upon the occurrence of a dendritic spike, a current, denoted 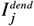, is generated in the dendrite and propagates to downstream regions of the cell. This current is characterized by its maximum amplitude *I*_*max*_, rise time *τ*_*rise*_, decay time *τ*_*decay*_, and delay Δ*t*^*s*^, and is defined as follows:

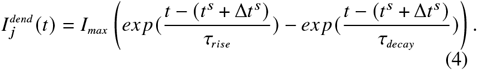

Here, *t*^*s*^ denotes the time at which dendritic spikes are generated. Note that we include a delay, Δ*t*^*s*^, to account for the time required for the dendrite to generate this current and for the current to propagate to the soma.

### Soma

The dendritic spike current is subject to perisomatic inhibition before being integrated at the soma. Consequently, we model the soma as integrating dendritic currents 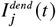 from nonAcD branches together with perisomatic inhibition, which is represented as a direct inhibitory current,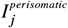. The resulting net excitatory current is then transmitted to the AIS, implemented as follows:

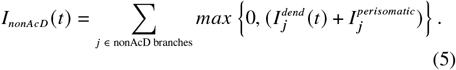

In nonAcD cells, *I*_*nonAcD*_ (*t*) is the only current reaching the AIS, whereas in AcD cells, an additional current from the AcD branch also contributes directly to the AIS input. The total positive current reaching the AIS, *I*^*inp*^ (*t*), is therefore given by:

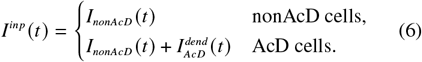

### Axon Initial Segment (AIS)

The membrane potential of the AIS determines whether an AP will be initiated. Let *V* ^*A*^ (*t*) represent the membrane potential of the AIS, which is characterized by a membrane resistance *R*_*A*_, a membrane capacitance *C*_*A*_, and a resting potential 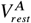. The membrane potential is calculated as follows:

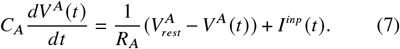

*I*^*inp*^ (*t*) denotes the total positive input current reaching the AIS from the soma and dendritic branches. When the membrane potential of the AIS exceeds the firing threshold 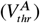, an action potential is elicited. The AIS then enters a refractory period, during which its membrane potential decays to the reset value 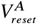 immediately after the spike (in this model, 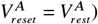.

### Synapses

A putative synapse between a dendritic branch *j* and a CA3 neuron *k* is represented by *θ*_*jk*_. This putative connection becomes a functional synapse with weight *w*_*jk*_ = *c*_*θ*_ *θ*_*jk*_ when its value exceeds zero. The dynamics of the synaptic weights are partially adapted from previous computational models [16, 22] and are defined as follows:

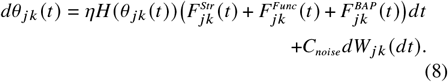

Here, *η* is the learning rate. The Heaviside function (*H*(·) ensures that the following plasticity terms only influence functional synapses (*θ*_*ki*_ (*t*) *>* 0).

### Structural Plasticity 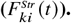

This component captures the dynamics of synaptic rewiring by imposing a structural limitation on how many functional synapses a dendritic branch can form. An illustration of this function is shown in Figure 3A.

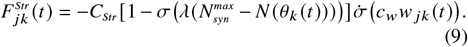

**Figure 3:**
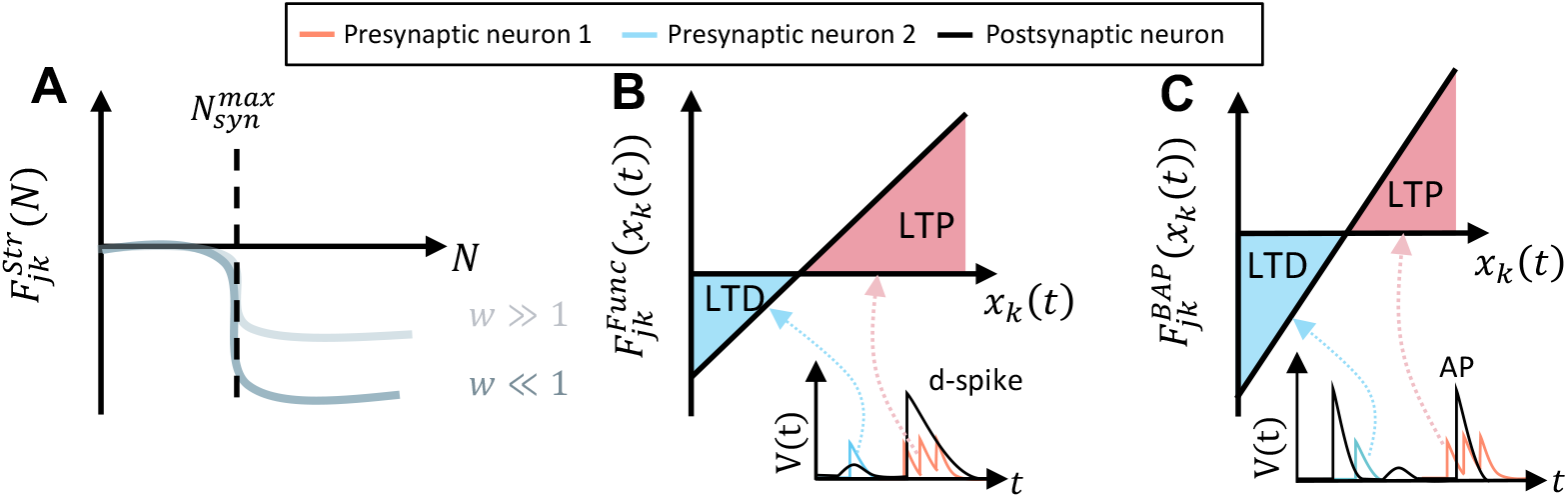
Synaptic plasticity. **A)** *Structural Plasticity*. This mechanism imposes a structural constraint on each dendritic branch, capping the number of synapses at a maximum value 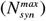. If this threshold is exceeded, all synapses on the branch are downscaled proportionally to their current weights. Consequently, strong synapses (*w* ≫ 1) are only slightly affected, whereas weaker ones (*w* ≪ 1) experience a more pronounced reduction. **B)** *Functional Plasticity*. When a dendritic spike is triggered within a branch, the strengths of its synapses are updated according to the activity level of their presynaptic neurons. Connections to highly active presynaptic neurons during the dendritic spike (red trace) are reinforced (LTP), whereas connections to less active presynaptic neurons (blue trace) are weakened (LTD). **C)** *BAP Plasticity*. When an AP is generated at the soma, it back-propagates into the dendritic tree, providing a global signal that modulates synaptic weights. This mechanism strengthens connections to presynaptic neurons that are highly active during the AP (red trace), while weakening connections to less active presynaptic neurons (blue trace). The dynamics of the synaptic weights are partially adapted from previous computational models [16, 22].

Here, the parameters *C*_*Str*_, *λ*, and *c*_*w*_ are positive scaling factors. The sigmoid *σ*(·) acts as a saturating term that constrains the number of synapses per branch *N* (*θ*_*k*_) to a maximum of 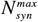. Experimental data has shown that the mean density of spines and average dendritic length of basal dendrites in nonAcD cells are 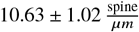 and 446.33 ± 168.13 *μm*, respectively [30]. Additionally, in AcD cells these parameters are reported as 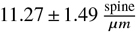 and 403.60 ± 285.40 *μm* for nonAcD branches, and 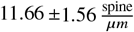 and 713.60 ± 371.50 *μm* for AcD branches [30]. These values result in 4747.61 ± 1839.79 spines per basal branch in nonAcD cells, and 4548.57 ± 3272.19 spines on nonAcD branches and 8320.58 ± 4472.45 spines on AcD branches of AcD cells. We maintain this ratio but scale down and round up the values for our simplified model. Therefore, we set the maximum number of synapses 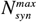 for AcD branches to 18 connections, and for nonAcD branches of both AcD and nonAcD cells to 10. Setting these values as the upper limit for the number of connections resulted in a number of connections that closely match experimental data, as shown in Figure 4.

**Figure 4:**
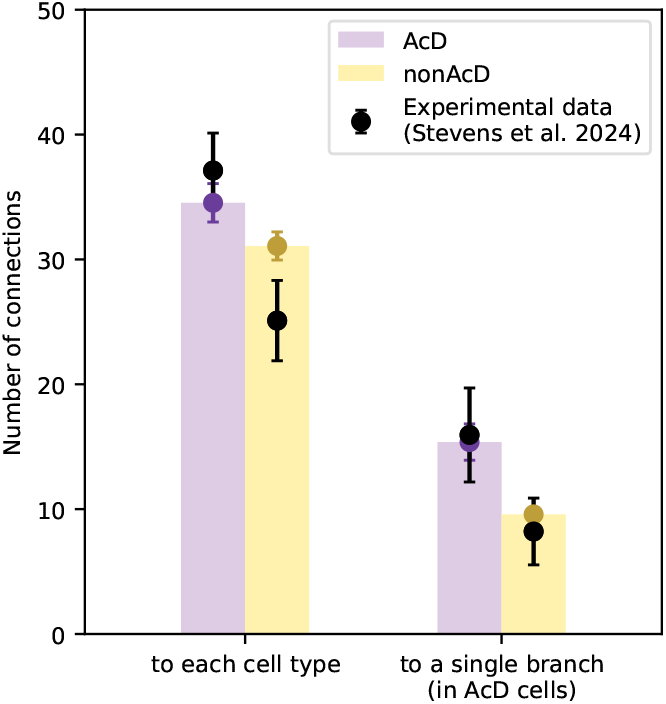
Number of synaptic connections to AcD and nonAcD cells and branches. The model was initialized with random connections, with the maximum number of synapses set to 10 for nonAcD branches in both cell types and 18 for AcD branches (based on calculations in the Structural Plasticity, 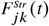, section). The network was then run until it reached equilibrium, after which the number of functional synapses on each branch was compared with spine–axon approximations (potential synapses) reported in Stevens et al. (2024) [30], showing strong agreement.

The derivative of the sigmoid,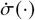, introduces a selective pruning effect: weaker synapses are penalized more strongly than stronger ones when *N*(*θ*_*k*_)exceeds 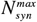. This behavior aligns with experimental evidence showing that larger spines (corresponding to stronger synapses) are more stable in vivo than smaller, weaker spines [15, 18, 23, 32].

Note that this term is always either zero or negative. Therefore, the structural plasticity term can only weaken the connections.

### Functional Plasticity 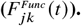

This component governs how synaptic weights are strengthened or weakened depending on the presynaptic neuron’s activity during dendritic spike events (see Figure 3B). The rule is expressed as:

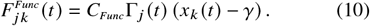

Here, *C*_*Func*_ is a scaling constant that determines the overall strength of functional plasticity, while *γ* specifies the critical activity level at which the rule switches from potentiation to depression (see Table 1). The function Γ_*j*_ (*t*) serves as a dendritic spike detector, taking a value of one during a spike and zero otherwise. The presynaptic activity trace of input neuron *k* is defined as 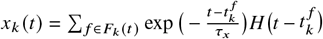 where *τ*_*x*_ is the time constant controlling the decay of presynaptic activity traces (see Table 1).

**Table 1.**
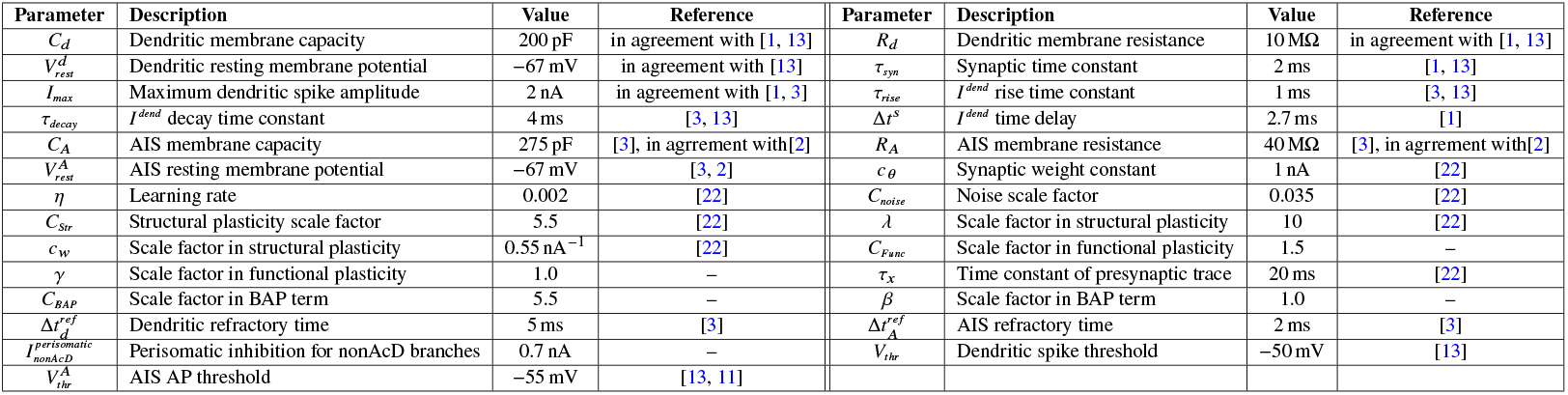
Model parameters.

In contrast to the structural plasticity, the functional plasticity can both strengthen and weaken the connections.

This depends on the activity of the CA3 input neuron *k*. If the neuron is highly active, i.e., *x*_*k*_ (*t*) *> γ*, the functional term will be positive, resulting in Long-Term Potentiation (LTP). Conversely, if the connected CA3 neuron has low activity, i.e., *x*_*k*_ (*t*) *< γ*, the functional term will be negative, leading to Long-Term Depression (LTD).

### Backpropagating Action Potential (BAP) Plasticity 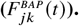

BAPs occur when an AP generated at the soma propagates back into the dendritic tree. This back-propagating current conveys a global signal to all dendritic branches, enabling the modification of synaptic connections. Although the current model does not explicitly simulate the back-propagating current, its effects on synaptic weight plasticity are incorporated. The implementation is as follows: whenever an AP is generated, all synaptic weights are updated based on their respective contributions to the AP stimulation. Specifically, if a presynaptic neuron exhibits high activity (i.e., *x*_*k*_ (*t*) *> β*) just before and during the AP, it is inferred that the corresponding synapse significantly contributed to the AP’s generation. As a result, such synapses are strengthened. Conversely, if the presynaptic neuron shows low activity (i.e., *x*_*k*_ (*t*) *< β*), indicating minimal or no contribution to the AP’s generation, the synapse is weakened (see Figure 3C).

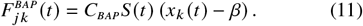

Here *C*_*BAP*_ is a positive scaling factor, and *β* defines threshold activity level required for an input neuron to contribute to action potential generation. *S* (*t*) represents the postsynaptic spike train:

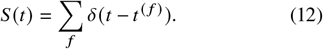

### Synaptic Noise

The stochastic aspect of the model arises from the final term in Equation 8. This term introduces the possibility of creating new functional synapses, as well as adding small random changes to the synaptic weights of existing functional synapses. In this formulation *dW*_*jk*_ (*dt*)

### Synaptic Noise

The stochastic aspect of the model arises from the final term in Equation 8. This term introduces the possibility of creating new functional synapses, as well as adding small random changes to the synaptic weights of existing functional synapses. In this formulation *dW*_*jk*_ (*dt*) corresponds to increments of a standard Wiener process, and the parameter *C*_*noise*_ sets the amplitude of the noise contribution.

## Results

We begin by showing that AcD and nonAcD cells adopt different learning mechanisms. Next, we demonstrate that the learned synaptic weights of AcD cells remain stable even during periods of strong inhibition. To achieve this, we developed a two-phase simulation framework: the *Exploration Phase*, modeling learning, and the *Ripple Phase*, assessing synaptic stability under inhibitory conditions.

### Exploration Phase

During spatial exploration in rodents, place-cell activity is strongly associated with theta oscillations, a prominent feature of hippocampal local field potentials, characterized by an 8*Hz* frequency. [7, 19, 20, 28, 29]. To simulate this rhythm in our model, CA3 cell assemblies were activated in random order, alternating between training and rest episodes of 60 ms each, yielding an oscillation period of 120 ms (≈8*Hz* frequency). During each training episode, a randomly selected assembly was activated as a Poisson spiking group at 100 Hz, while background activity from neurons in other assemblies was maintained at 1 Hz. In the subsequent rest episode, all input neurons were activated at 1 Hz. An illustration of this protocol is shown in Figure 5A. Figure 5B and C illustrate examples of synaptic weight sum evolution during the learning phase for an AcD cell and a nonAcD cell, respectively. Note that in the AcD cell (Figure 5B), branch 1—the axon-carrying dendrite—exhibits higher maximal synaptic weights compared to the nonAcD branches. This effect arises from the greater number of synaptic sites on the AcD branch, which is a consequence of its extended length.

**Figure 5:**
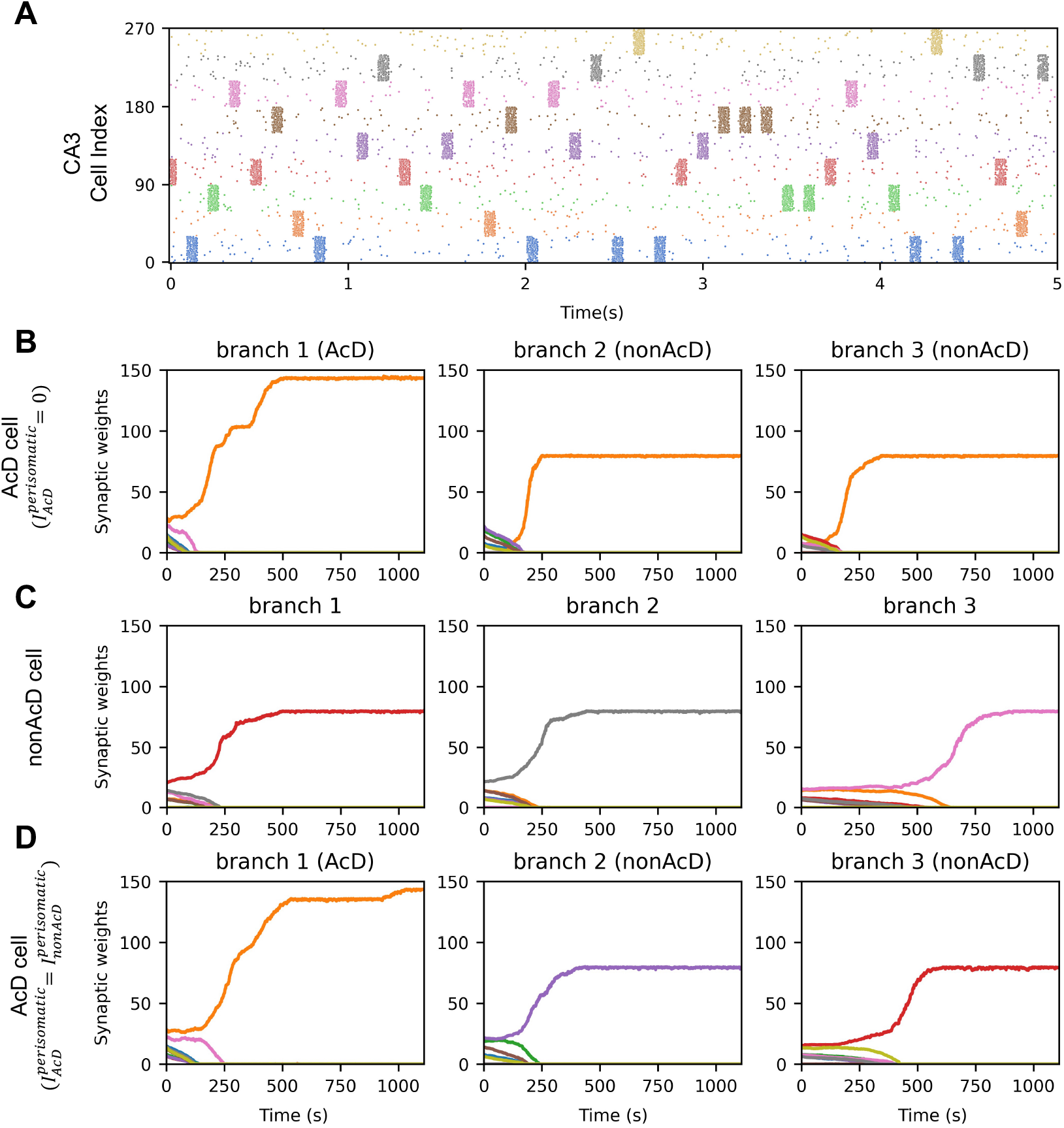
Exploration Phase. **A)** *Simulation Protocol:* The simulation consists of alternating 60 ms episodes of activity and rest, corresponding to a frequency of approximately 8 Hz (theta rhythm). During each active episode, one assembly (shown in a distinct color) is randomly selected and activated at 100 Hz, while the remaining assemblies are active at 1 Hz to simulate background noise. During rest episodes, all assemblies are active only at the background noise level (1 Hz). **B)** *AcD Cell:* Synaptic weight dynamics for the three dendritic branches of an AcD cell during the exploration phase. Each curve shows the summed synaptic weight of connections between a branch and a specific assembly. The AcD branch reaches higher maximum weights than the nonAcD branches, reflecting its greater number of synaptic sites. Starting from random CA3–CA1 connections, the privileged AcD pathway dictates which assembly the other branches learn, resulting in a single-modal neuron. **C)** *NonAcD Cell:* Same as panel B, but for a nonAcD cell. In nonAcD cells, all dendritic branches are equivalent, with no privileged pathways, so each branch learns a different assembly, resulting in a multi-modal neuron. **D)** *AcD Cell with Perisomatic Inhibition on AcD Branch:* Same as panels B and C, but for an AcD cell in which the AcD branch receives perisomatic inhibition and lacks its privileged input pathway. Normally, bypassing perisomatic inhibition enhances AP generation, amplifying the BAP signal and plasticity, which drives other branches to learn the same assembly. With the privileged pathway disabled, this BAP-mediated effect is removed, preventing the AcD cell from becoming single-modal.

At the onset of the learning phase, synaptic connections between assemblies and dendritic branches were random. After 1000 s of learning, each dendritic branch had selectively learned to respond to a specific assembly. In AcD cells, the privileged pathway of the AcD branch played a central role, guiding the learning process of the other branches and determining which assembly they connected to. This mechanism produced a **single-modalized** cell, in which all branches learn to respond to a unique assembly. In contrast, in nonAcD cells, where no branch held precedence, branches learned different assemblies, resulting in a **multi-modalized** cell. A schematic representation of a single-modalized AcD cell and a multi-modalized nonAcD cell is shown in Figure 6. Moreover, in the simulated network of 50 AcD cells and 50 nonAcD cells, all AcD cells became single-modalized, whereas the majority of nonAcD cells were multi-modalized (Figure 7).

**Figure 6:**
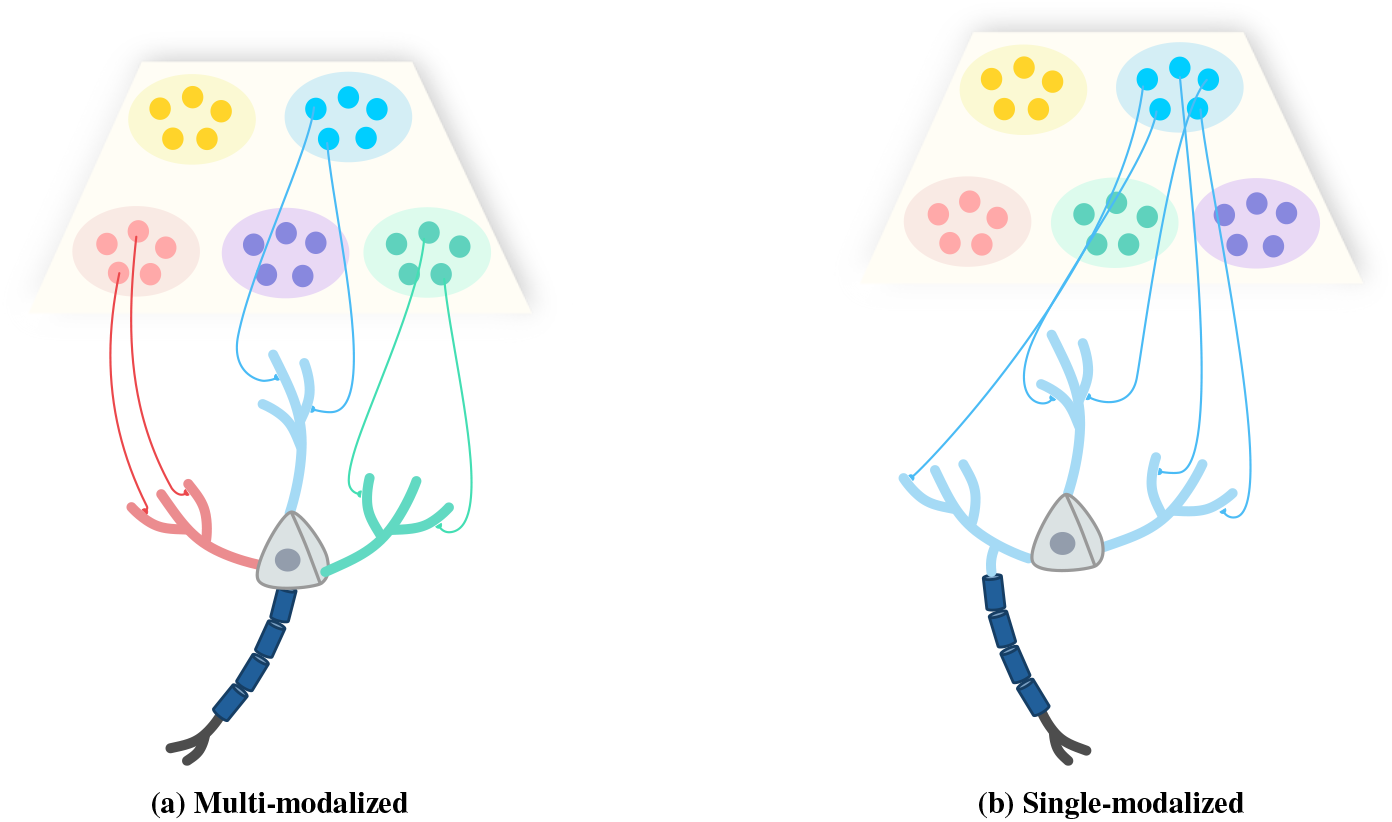
Multi-modalized vs. Single-modalized. Illustration of a multi-modal nonAcD cell (a) and a single-modal AcD cell (b). Colored boxes denote distinct CA3 neuron assemblies. A dendritic branch learns an input assembly if it forms a sufficient number of strong connections with it. In the multi-modal cell, each branch learns a different assembly, whereas in the single-modal cell, all branches converge on the same assembly.

**Figure 7:**
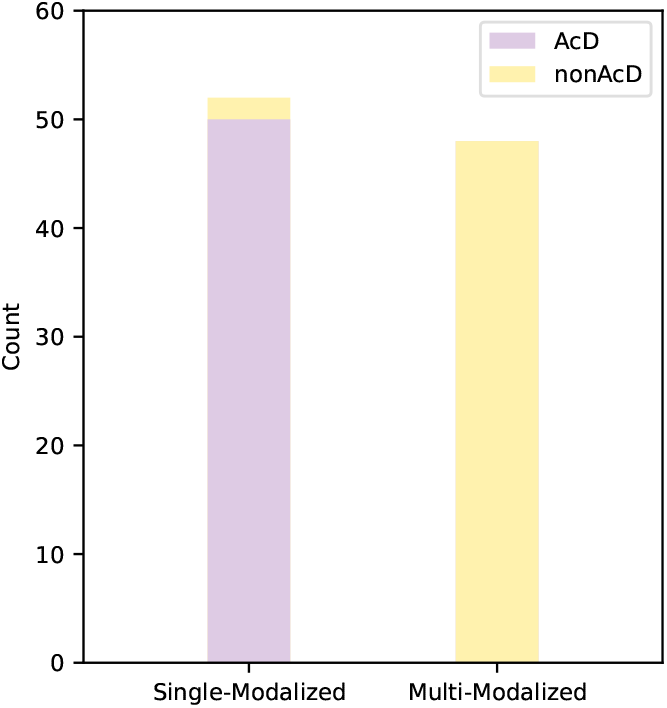
Single-modalized vs. Multi-modalized Population. In this simulated network of 50 AcD and 50 nonAcD cells, at the end of the exploration phase, all AcD cells are single-modalized, whereas the majority of nonAcD cells exhibit multi-modalized learning mechanism.

To evaluate the effect of the privileged input pathway of AcD branches—which bypasses perisomatic inhibition—we performed an additional simulation in which AcD branches were subjected to the same perisomatic inhibition as nonAcD branches 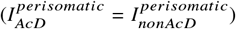. In this setting, the sole difference between AcD and nonAcD branches was the larger number of synaptic connections on AcD branches, attributable to their greater length. The corresponding results are presented in Figure 5D. These results demonstrate that, in the absence of the privileged input pathway of AcD branches, both cell types adopt a multi-modal organization. We conclude that the lower AP generation threshold for inputs to AcD branches—arising from their ability to bypass perisomatic inhibition—enhances AP generation during activity of the assembly learned by the AcD branch. This heightened activity amplifies BAP-mediated plasticity in nonAcD branches, thereby compelling them to learn the same assembly. Consequently, the bypassing of perisomatic inhibition in AcD branches establishes a distinct learning mechanism, yielding a single-modal learning pattern in AcD cells, in contrast to the multi-modal pattern observed in nonAcD cells.

### Ripple Phase

Hippocampal sharp-wave ripple complexes (SWRs) are brief, high-frequency oscillatory events that occur during slow-wave sleep and passive wakeful states, such as eating, drinking, grooming, and quiet wakefulness, with a frequency of occurrence ranging from 0.01 to 2 Hz [5, 6, 25]. Sharp waves are triggered by the coordinated bursting of CA3 neuron ensembles, resulting in extensive, non-rhythmic depolarization of the apical dendrites of CA1 pyramidal cells within the stratum radiatum [8]. The dense interconnectivity of CA1 interneurons result in their extensive recruitment and highly synchronized firing at frequencies of 150–250 Hz during sharp waves [4, 24]. Evidence from some studies suggests that perisomatic interneurons are key regulators in determining the specific assembly of CA3 neurons involved in each SWR [10]. The interplay between oscillatory inhibition of interneurons and strong excitation from CA3 produces rapid field oscillations in the CA1 pyramidal layer, referred to as ripples [6, 37]. Experimental evidence suggests that disrupting SWRs during the sleep phase impairs memory consolidation [8, 9]. Furthermore, awake ripples contribute to memory consolidation by serving as a memory tag, identifying specific experiences for targeted consolidation during subsequent sleep [36]. Recent experimental findings suggest that during ripple events in awake mice, AcD cells exhibit significantly higher firing probabilities and frequencies compared to nonAcD cells [17]. Building on this observation, we examine how the elevated firing probability of AcD cells during periods of high inhibition, such as ripple events, influences the stability of learned synaptic weights. To explore this, we simulated the 2 Hz event frequency of the ripples by structuring the phase into 500 ms cycles, each consisting of 100 ms of CA3 activity that induces CA1 cell depolarization, followed by 400 ms of rest. Figure 8A provides an illustration of this protocol. In each episode of CA3 activity, a random assembly is selected, and fires at a rate of 100 Hz, with background noise maintained at 1 Hz. During CA3 neuronal activity, an inhibitory current with a frequency of 150 Hz, denoted as 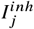, is applied to each dendritic branch of the CA1 pyramidal cells. This current simulates the activity of CA1 interneurons and their synchronized high-frequency activity during ripple events [4, 24]. An example of the weight evolution during the ripple phase for a pair of representative AcD and nonAcD cells is shown in Figure 8B and C, respectively. As demonstrated, the learned synaptic weights across all branches of the AcD cell remain stable (Figure 8B), whereas the synaptic weights of the nonAcD cell (Figure 8C) decay during this period of high inhibition. To investigate the role of the privileged input pathway in synaptic weight stability, we conducted a simulation with equivalent perisomatic inhibition for both AcD and nonAcD branches. The results, shown in Figure 8D, reveal that in this scenario, only the synaptic connections on the AcD branches remain stable. This occurs because, even during periods of high inhibition due to interneurons activity, the increased dendritic spike propensity in the AcD branches leads to a greater number of dendritic spikes compared to the nonAcD branches. As a result, the functional term in synaptic plasticity contributes more positively, aiding in the consolidation of synaptic weights on the AcD branches, regardless of whether perisomatic inhibition is bypassed.

**Figure 8:**
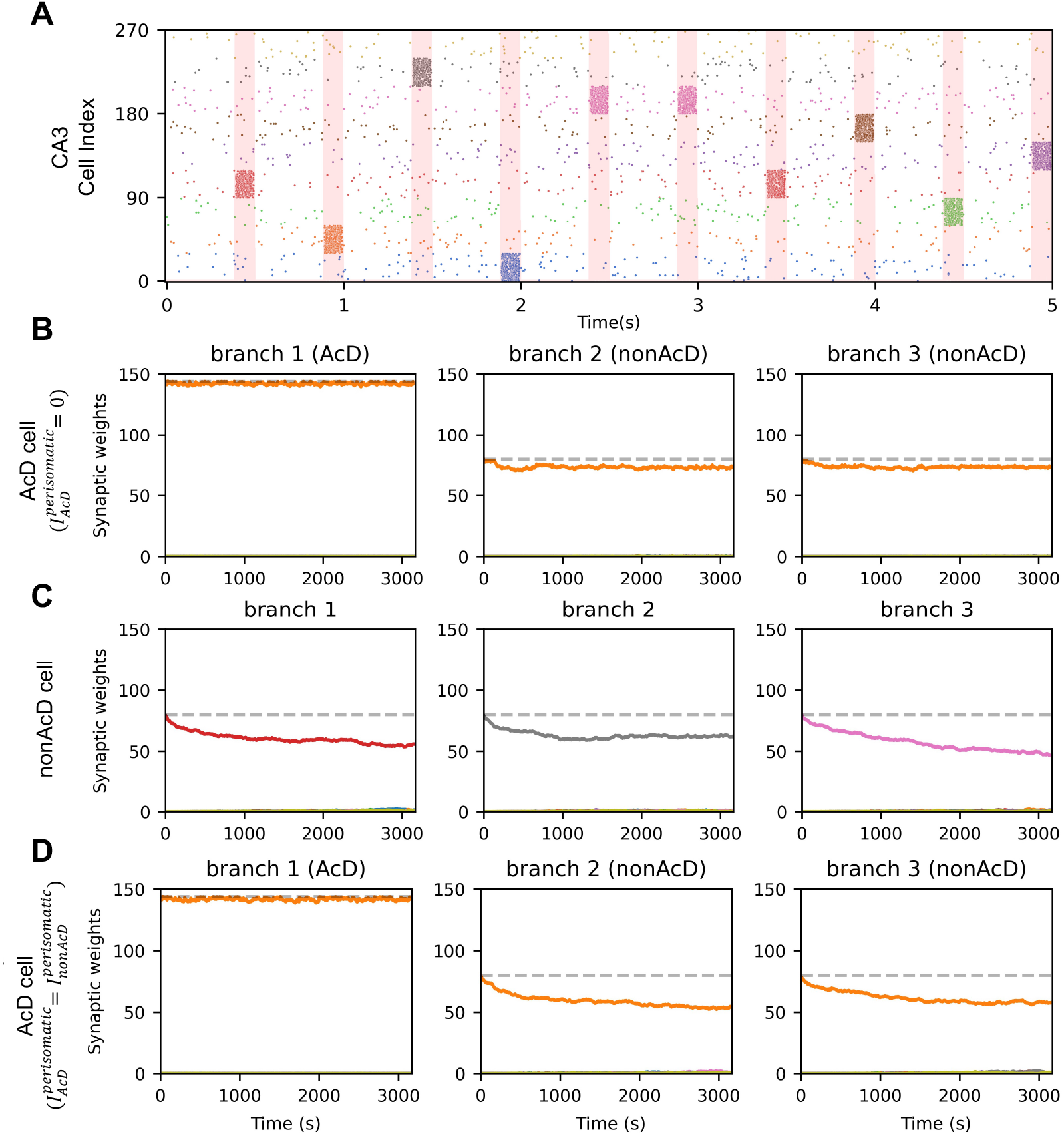
Exploration Phase. **A)** *Simulation Protocol:* To simulate the 2 Hz SWR event frequency, this phase is divided into 500 ms cycles, consisting of 100 ms of activity and 400 ms of rest. During the active episodes, one random assembly is activated at 100 Hz frequency, while others are active at the background noise level (1 Hz). Additionally, to mimic the synchronized high-frequency activity of interneurons, a 150 Hz inhibitory current is applied to each dendritic compartment during the active periods (indicated by the shaded red regions). During the longer rest periods, all CA3 assemblies are active at the background noise level, and no inhibitory current is applied to the branches. **B)** *AcD Cell:* A sample synaptic weight evolution for all three dendritic branches of an AcD cell is plotted during the ripple phase. Each curve represent the weight sum of all the synaptic connections between that branch and a specific assembly. Note that, the maximum synaptic weight for the AcD branch is higher than that of the nonAcD branches, reflecting the increased capacity of AcD branches to form synaptic connections, due to their greater number of synaptic sites. This phase begins with initialized learned synaptic weights. During the ripple phase, the synaptic weights on all branches of the AcD cell remain stable and preserved. **C)** *nonAcD Cell:* Same as B, but for a sample nonAcD cell. The synaptic weights on all branches gradually decay. **D)** *AcD Cell with Perisomatic Inhibition on AcD Branch:* This simulation applies the ripple-phase protocol while blocking the privileged input pathway of AcD cells by applying perisomatic inhibition to the AcD branch at the same level as the nonAcD branches (*I*^perisomatic^AcD = *I*^perisomatic^nonAcD). Under these conditions, only the synaptic connections on the AcD branch remain stable, whereas the connections on the two nonAcD branches decay. These findings highlight the critical role of the privileged input pathway of the AcD branch in maintaining the stability of synaptic weights in AcD cells.

Additionally, in the simulation with perisomatic inhibition bypassing, these dendritic spikes on the AcD branches facilitate AP generation, resulting in BAPs in the nonAcD branches of the AcD cell. Consequently, the synaptic connections on the nonAcD branches receive positive reinforcement from the BAP term, despite their limited ability to generate dendritic spikes due to high inhibitory currents. This results in the consolidation and stability of synaptic weights on both AcD and nonAcD branches. In contrast, the lack of sufficient dendritic spikes and APs in nonAcD cells leads to synaptic weight decay and instability.

## Discussion

In this study, we introduced a simplified CA3-CA1 network model to investigate how the privileged input pathway of AcD cells influences their ability to learn and stabilize information. Our findings indicate that AcD and nonAcD cells, despite sharing similar morphological features, exhibit distinct learning behaviors. AcD cells tend to become single-modalized, meaning that all their dendritic branches predominantly learn the same input assembly. In contrast, nonAcD cells are more likely to become multi-modalized, where each branch learns distinct input assemblies. Furthermore, the synaptic connections on AcD cells demonstrate greater stability during high-inhibition phases, such as ripple events. This observation is consistent with experimental data showing that CA1 pyramidal cells in the superficial layers, where AcD cells are more prevalent, exhibit enhanced stability compared to pyramidal cells in the deeper layers, which predominantly consist of nonAcD cells [14, 31, 33]. Another study has shown that CA1 place cells in the superficial sublayer are more active in cue-poor environments, relying predominantly on a firing rate code driven by intra-hippocampal inputs [27]. In contrast, CA1 place cells in the deep sublayer are more active in cue-rich environments and utilize a phase code driven by entorhinal inputs [27]. Consistent with this, our model suggests that AcD cells operate under a single-modalized mechanism, where their activity is driven by a single CA3 assembly. Consequently, in cue-poor environments, which are characterized by fewer landmarks and fewer active CA3 assemblies, AcD cells—predominantly located in the superficial layer of CA1—remain capable of generating APs. In contrast, nonAcD cells, which employ a multi-modalized mechanism, require input from multiple active assemblies to generate APs. As a result, nonAcD cells, which are more prevalent in the deep layers, are less active in cue-poor environments. Furthermore, the multi-modalized mechanism of nonAcD cells enables them to encode distinct landmarks across their branches, facilitating the representation of more complex, cue-rich environments.

A recent study has shown that awake ripples serve as memory tags, marking specific experiences for consolidation during sleep [36]. Our results indicate that during periods of high inhibition, the information encoded in AcD cells remains more stable than that in nonAcD cells. This suggests that AcD cells are better suited for storing critical information that is prioritized for consolidation during sleep-associated SWRs.

These findings highlight the distinct roles of AcD and nonAcD cells in spatial coding and memory consolidation. Single-modalized AcD cells, with their increased stability and specialization, are likely optimized for encoding a single spatial or contextual feature. This corresponds to place cells with highly specific and robust firing fields, which may be essential for tasks requiring high reliability and minimal interference, such as recalling a specific landmark or repeatedly navigating to a single destination. In contrast, the multi-modalized nature of nonAcD cells reflects their flexibility in encoding diverse or complex environmental information. These cells likely play a critical role in forming distributed place fields, enabling them to represent multiple spatial locations or experiences. This makes nonAcD cells vital for tasks that require integration of multiple features, such as navigating a maze with multiple landmarks or associating various contextual cues with specific outcomes.

In summary, our findings suggest that, despite similarities in morphology and EPSC kinetics between AcD and nonAcD cells [17, 30, 31], their functional roles in learning, synaptic weights stability, and consolidation diverge substantially. AcD cells prioritize specialization and stability, making them particularly useful in environments with minimal external cues. NonAcD cells, in contrast, provide the flexibility to encode and integrate information in environments rich with diverse features. Together, these distinct coding mechanisms ensure that the hippocampus can effectively navigate and adapt across heterogeneous spatial contexts.

